# Ventilation does not affect influenza virus transmission efficiency in a ferret playpen setup

**DOI:** 10.1101/2023.12.26.573248

**Authors:** Nicole C. Rockey, Valerie Le Sage, Meredith Shephard, Nahara Vargas-Maldonado, Andrea J. French, Sydney Walter, Lucas M. Ferreri, Katie E. Holmes, David VanInsberghe, Herek Clack, Aaron J. Prussin, Anice C. Lowen, Linsey C. Marr, Seema S. Lakdawala

## Abstract

Sustained community spread of influenza viruses relies on efficient person-to-person transmission. Current experimental transmission systems do not mimic environmental conditions (e.g., air exchange rates, flow patterns), host behaviors or exposure durations relevant to real-world settings. Therefore, results from these traditional systems may not be representative of influenza virus transmission in humans. To address this pitfall, we developed a modified, more realistic transmission setup and used it to investigate the impact of ventilation rates on transmission in a close-range, play-based scenario. In this setup, four immunologically naïve recipient ferrets were exposed to a donor ferret infected with a genetically barcoded 2009 H1N1 virus (H1N1pdm09) for four hours. The ferrets interacted in a shared space that included toys, similar to a child care setting. The transmission efficiency was determined under conditions of low and high ventilation rates; air exchange rates of ∼ 1.3 hr^-1^ and 23 hr^-1^, respectively. Despite the large difference in ventilation rate, transmission efficiency was the same, 50% in two independent replicate studies. The presence of infectious virus or viral RNA on surfaces and in air throughout the exposure area was similar regardless of ventilation rate. While high viral genetic diversity in donor ferret nasal washes was maintained during infection, recipient ferret nasal washes displayed low diversity, revealing a narrow transmission bottleneck regardless of ventilation rate. Our findings indicate that in exposures characterized by frequent close-range, play-based interactions and the presence of fomites, ventilation does not significantly impact transmission efficiency.

**Significance:** Improved ventilation in building has the potential to reduce transmission of respiratory viruses, but its effect in different settings is not well understood. We developed a novel system to study influenza virus transmission in the ferret animal model in an environment that mimics a child care center. We demonstrate that increased ventilation is not effective at disrupting transmission in this setting, suggesting that transmission occurs mainly at close-range or via fomites. Multiple interventions are needed to reduce the spread of influenza virus in this type of setting.

## Introduction

Seasonal and pandemic influenza viruses cause tens of thousands of deaths each year in the US (1). To circulate successfully in the human population, viruses must sustain human-to-human spread. Influenza virus transmission occurs via numerous routes, including direct and indirect (e.g., fomites) contact, inhalation of aerosols, and spray of large droplets (2–6). While all modes of transmission may occur during an exposure event, the relative contribution of each of these routes varies depending on host, environmental, and viral factors (3).

Certain settings are linked with elevated rates of transmission, including households and child care settings (7–10). In these situations, individuals interact in close proximity for extended periods of time (e.g., hours). Understanding how influenza virus transmission occurs and what interventions may reduce spread in these settings is critical, but conclusive data is lacking. Randomized control trials and observational epidemiological studies in combination with modeling efforts have expanded knowledge in this area, and while some find marginal associations of hand hygiene and face mask use with lower levels of illness, most studies do not detect a statistical difference (11–19). Unfortunately, these studies have not decisively identified effective prevention strategies or predominant transmission routes because of confounding factors or limited adherence to interventions.

To fill this knowledge gap, controlled human transmission studies have been suggested (20). To date, two human influenza transmission studies have been conducted to mimic exposures in close living spaces (21, 22). The first proof-of-concept pilot resulted in transmission to three of 12 recipients (22), while the higher-powered study only observed transmission to one of 75 recipients (21). The expense of human challenge research and difficulties in consistently attaining influenza virus transmission have constrained this work.

More commonly, animal models of influenza virus transmission, and in particular the ferret model, are used to assess transmission dynamics of influenza viruses (23). Results of prior studies may not be representative of transmission in real-world contexts since the experimental setups typically use directional airflow, elevated ventilation, lengthy exposure durations, and/or restricted donor-to-recipient interactions (24–32). While some studies have used shortened exposure durations to shift towards more realistic transmission systems (33, 34), a number of features still do not recapitulate everyday interactions among humans.

To reduce respiratory virus spread in high-risk settings, non-pharmaceutical interventions are often recommended (35, 36). Common environmental controls include barriers, physical distancing, masking, filtration, ventilation, and humidification. These strategies aim to decrease exposure to infectious virus in the air. Increasing ventilation, which reduces concentrations of contaminants in air and subsequent deposition of virus on surfaces, has been suggested to reduce influenza virus transmission rates (37–41). However, these studies contain significant limitations: modeling components rely on unvalidated assumptions and observational studies often include uncontrolled variables such that causality cannot be firmly established. Thus, an experimental setting that mimics real-world transmission scenarios is critical for studying how viruses transmit and assessing the impact of recommended engineering controls, such as ventilation, on virus spread.

Transmission to a new host often vastly changes the genetic composition of a viral population. This effect has important implications for viral evolution and can potentially also offer insight into the physical and biological processes that shape the transmission event. Indeed, previous animal studies evaluating transmission bottlenecks provided evidence that the route of infection influences the diversity transferred between hosts (42, 43). Nevertheless, it remains unclear how environmental conditions and host behavior impact the transfer of viral diversity between hosts during a transmission event.

In this study, we describe a novel exposure environment for studying influenza virus transmission among ferrets that more accurately recapitulates transmission in a scenario involving frequent close-range interactions among humans over several hours, such as those observed in child care settings. This system was used to assess how ventilation affects environmental contamination and transmission in play-based exposures. To increase the resolution of viral detection in recipient ferrets, we used a barcoded virus population. Our results indicate that, in a play-based scenario, increasing ventilation by a factor of 18 does not affect transmission efficiency or the number of distinct viral variants that establish infection. This work has important implications for informing the value of engineering controls in indoor environments. Our findings also highlight the role that exposure setting and associated behaviors may play in driving transmission, even in the presence of engineering controls like ventilation.

## Results

### Play-based system to study influenza virus transmission in ferrets

To examine influenza virus transmission efficiency in a more realistic setting, we developed a new ferret model system that allows for environmental conditions and behavior like those in a child care setting. Children in daycare facilities exhibit a wide range of behaviors and interactions, including close interactions with one another (e.g., touching, hugging) and exploratory behaviors involving touching and sharing toys or other objects. To simulate a child care setting under controlled experimental conditions, we used ferrets because their behaviors recapitulate many of those exhibited by children. Ferrets are curious animals that explore and play with toys and other surfaces through touch and bite behavior. They are also social animals and display considerable close-range behaviors, including chasing, biting, nuzzling, and cuddling (44).

In our ferret transmission setup, recipient ferrets were housed with a single infected donor ferret in a defined area (∼ 3.5 m^2^) for a single exposure period (see Fig.1, Methods, Movie S1, and Movie S2 for more details). This system allows for all modes of transmission: contact (including from toys and other objects), aerosol, and droplet spray. In each transmission experiment (Fig. 1A), the donor ferret was intranasally infected with 10^6^ TCID_50_ of barcoded H1Npdm09 virus, placed in the exposure area at one day post-infection and allowed to interact with four naïve recipient ferrets for four continuous hours. Following the exposure period, all ferrets were individually housed. Nasal washes and oral swabs were collected to evaluate viral shedding and seroconversion of the recipients was determined on days 14 and 21 post donor infection (Fig. 1A).

**Figure 1.**
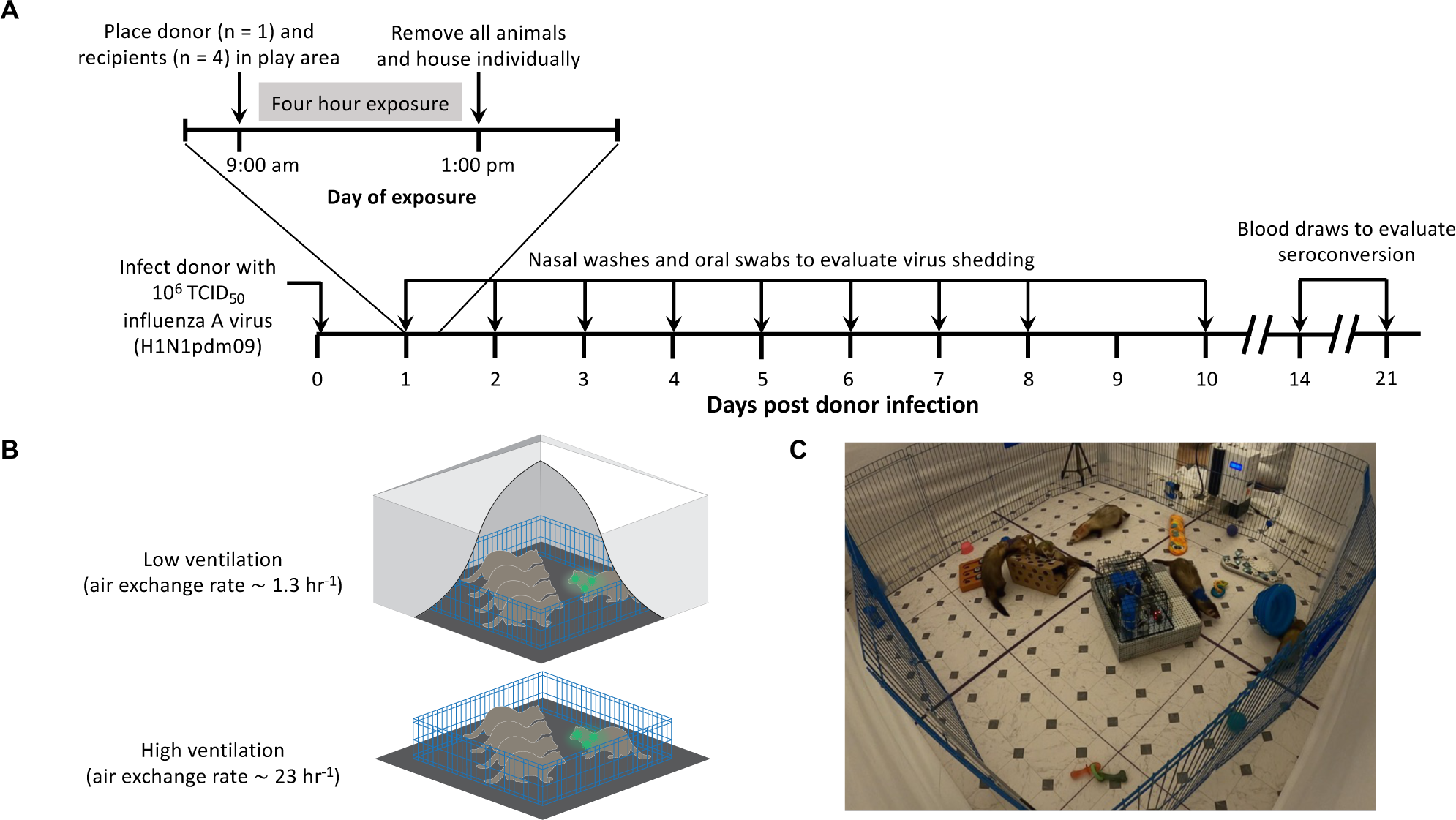
Overview of the close-contact, play-based ferret transmission setup. (A) Timeline of activities associated with the transmission experiments, (B) schematics of the transmission system for low and high ventilation settings, and (C) image of a transmission exposure experiment.

Our system enabled us to modulate ventilation conditions to a low air exchange rate of ∼1.3 air changes per hour, representative of those found in schools and child care facilities (45, 46), or a high rate of ∼23 air changes per hour (Fig. 1B, Movies S1 and S2), exceeding ventilation rates that are recommended for healthcare facilities (47). This allowed us to assess the impact of elevated ventilation rate on transmission. The low ventilation condition was achieved by enclosing the transmission system within a tent. The air exchange rate in the tent was experimentally confirmed using the CO_2_ decay method (see Methods, Fig. S1). The elevated ventilation condition was attained by removing the tent. In this case, the air exchange rate was equal to that of the room in the animal facility and was calculated based on mechanical ventilation parameters (see Methods). Two independent transmission studies were performed for each ventilation condition.

For each transmission experiment, toys, food, and water were placed throughout the space to encourage ferret-object interactions (Fig. 1C). Additionally, several different types of aerosol collection devices, namely National Institute for Occupational Safety and Health (NIOSH) bioaerosol cyclone samplers, gelatin cassette samplers, and liquid Spot samplers, were placed within the exposure space to measure aerosolized infectious virus and viral RNA (Fig. 2A). To assess fomite contamination, toys and other surfaces were swabbed post-exposure for infectious virus and viral RNA. CO_2_ levels, which are indicative of exhaled breath, were measured using Aranet4 sensors and were higher in the low ventilation condition, as expected (Fig. S2). These sensors were also used to monitor relative humidity and temperature at different locations within the exposure area (Fig. S3). A remote-controlled battery-powered robot collected aerosols at close-range to the animals using NIOSH bioaerosol cyclone samplers and a gelatin cassette sampler; an Aranet4 sensor housed within the robot measured humidity, temperature, and CO_2_ (Fig. 1C and Fig. 2A).

**Figure 2.**
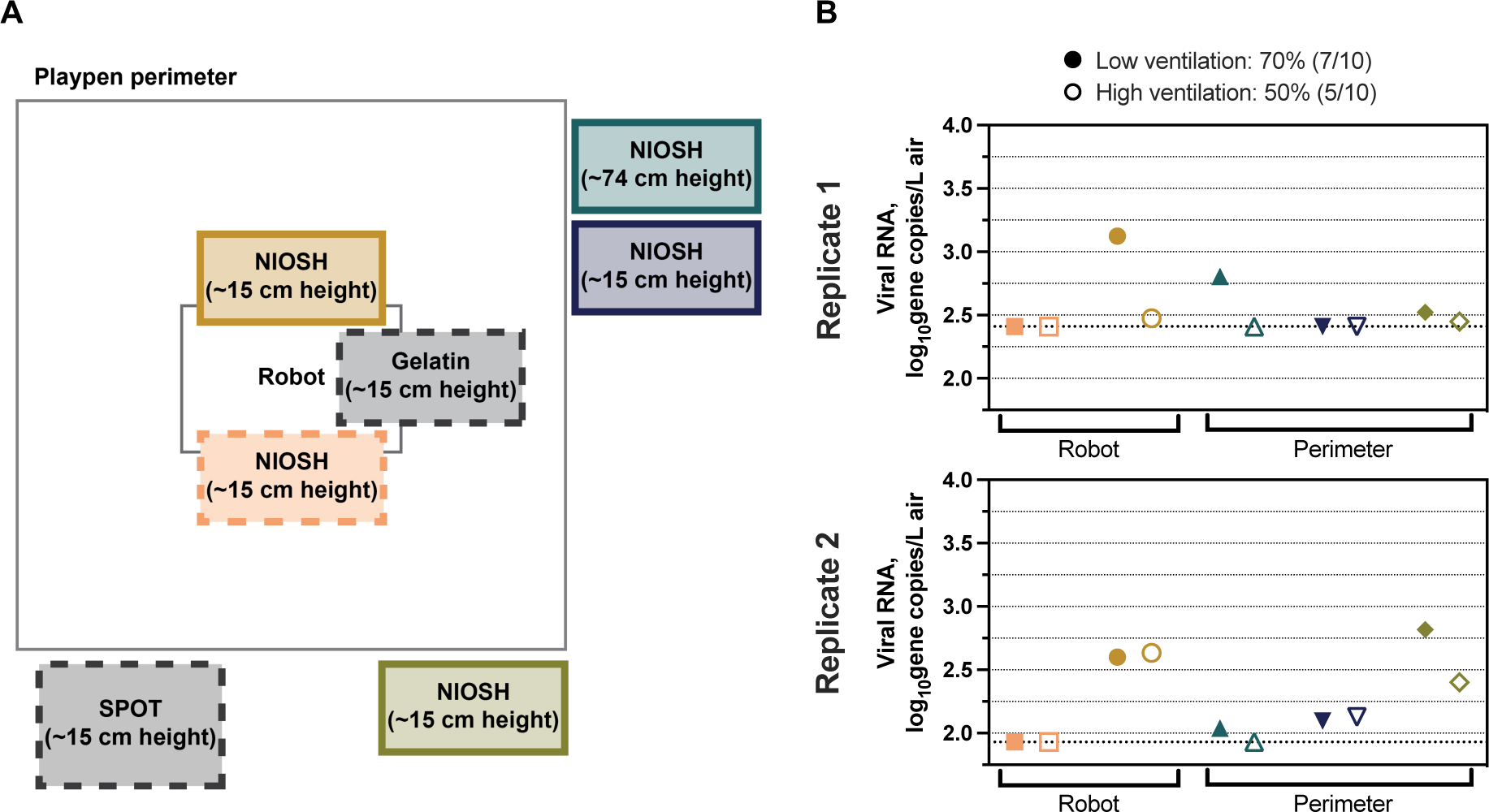
Ventilation setting did not significantly impact viral RNA concentrations detected in aerosols. (A) Schematic of air sampler locations, including the Spot Sampler aerosol particle collector (SPOT), gelatin cassette sampler (Gelatin), and NIOSH bioaerosol cyclone samplers (NIOSH). Dashed outlines indicate samplers used to detect both infectious virus and viral RNA, and solid outlines indicate samplers used to detect only viral RNA. (B) Total influenza virus RNA concentrations collected from each NIOSH bioaerosol cyclone sampler during two independent experiments with low (closed symbols) or high (open symbols) ventilation. All three size fractions of the NIOSH bioaerosol cyclone sampler were analyzed for presence of viral RNA separately, but are shown here as the sum of all fractions. The color of symbols in (B) correspond to the NIOSH bioaerosol cyclone sampler shown in (A). Dashed lines indicate the limit of detection. NIOSH samplers, SPOT sampler, and gelatin cassette sampler were each run for the entire four-hour exposure at flow rates of 3.5, 1.4, and 1.5 L/min, respectively. No infectious virus was detected from any air samplers, and no viral RNA was detected in the gelatin cassette or Spot Sampler.

### Viral RNA concentrations were not significantly lower when ventilation was increased

Ventilation is expected to reduce the concentrations of virus-containing aerosols within a space. To assess the impact of ventilation on influenza virus-containing aerosols, we compared the concentration of viral RNA in air from exposures with low and high ventilation rates (Fig. 2A). We did not detect infectious virus in samples collected from any of three air-sampling devices (data not shown), namely a Spot Sampler condensation particle collector, a NIOSH bioaerosol cyclone sampler, and a gelatin cassette sampler, even though infectious virus was detected from donor ferret aerosol expulsions sampled directly into the Spot Sampler (Fig. S4 and Methods). Although no infectious influenza virus was detected from the air in the experimental enclosure, viral RNA was captured using the NIOSH bioaerosol cyclone samplers in both the low and high ventilation settings (Fig. 2B). No viral RNA was detected from the gelatin cassette or Spot Sampler (data not shown). The proportion of positive aerosol samples and the concentrations of viral RNA detected were not significantly different between the low and high ventilation experiments. Across two independent experiments, 50% (5/10) of samplers were positive in the high ventilation setting, while 70% (7/10) of samplers were positive in the low ventilation setting. Concentrations of viral RNA sampled using the NIOSH bioaerosol cyclone samplers in the high ventilation experiment were slightly lower, on average 2.3 log_10_ gene copies/L air, than in the low ventilation experiment, at 2.5 log_10_ gene copies/L air, although this difference was not significant (Mann-Whitney test, p = 0.31). Therefore, increased ventilation did not appear to affect the amount of viral RNA in air sampled around the ferrets during the exposure period.

### Surfaces in the transmission enclosures contained infectious virus, regardless of ventilation rate

Our transmission setup also provided the opportunity for ferrets to interact with various toys and objects, similar to what may occur in a child care setting. Given the potential for influenza virus shedding and subsequent contamination of fomites during each exposure, we sampled surfaces and assessed the distribution and concentration of both infectious influenza virus and viral RNA (Fig. 3A). The proportion of surface samples positive for infectious influenza virus was the same across both replicate sets of experiments, with surface positivity in 13.5% of samples (7/52) in either ventilation setting (Fig. 3B). Concentrations were statistically similar across the distinct ventilation settings (Mann-Whitney test, p = 0.88), with a mean of 0.61 log_10_ TCID_50_/swab in the low ventilation setting and 0.62 log_10_ TCID_50_/swab in the high ventilation setting.

**Figure 3.**
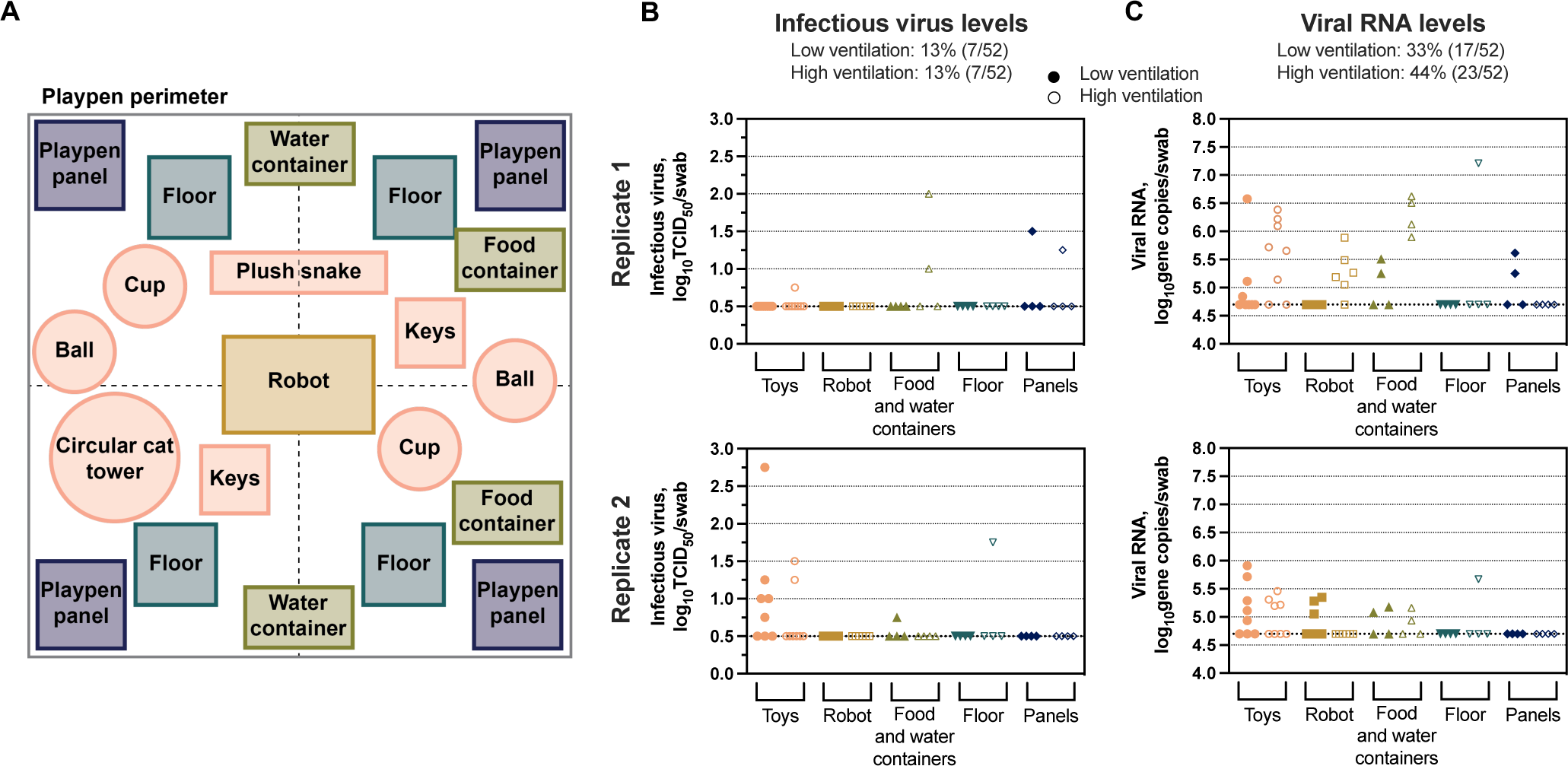
Similar influenza virus concentrations from surfaces sampled following low and high ventilation transmission exposures. (A) Schematic of all surfaces sampled following each transmission exposure. (B) Amount of infectious influenza virus detected on each surface grouped by object type for both replicates. (C) Total concentration of viral RNA per swab detected from surface samples group by object type during two independent experiments with low (closed symbols) or high (open symbols) ventilation. The color of symbols in (B) and (C) correspond to the samples of the same color shown in (A). Dashed lines indicate the limit of detection.

Detection of viral RNA on surfaces followed the same trend as that of infectious virus, yielding similar concentrations across the low and high ventilation experiments (Mann-Whitney test, p = 0.09; low ventilation mean = 4.9 log_10_ gene copies/swab, high ventilation mean = 5.1 log_10_ gene copies/swab; Fig. 3C). As expected, viral RNA positivity was higher than infectious virus positivity, with 33% (17/52) and 44% (23/52) of surface samples being positive for viral RNA in the low and high ventilation exposures, respectively. Together, these results provide evidence that ventilation did not significantly impact the extent of surface contamination that occurred due to viral shedding from an infected ferret.

### Increased ventilation did not reduce transmission in a play-based setting

Ultimately, we developed this transmission system to directly evaluate whether ventilation would affect the efficiency of influenza virus spread in a play-based setting. In both low and high ventilation settings, four of eight (50%) of recipients shed virus beginning at least one day post-exposure (Fig. 4, Fig. S5). Seroconversion of recipient ferrets post-exposure confirmed these findings (Table S1). Of note, the onset of viral shedding in the nasal washes of infected recipients from the high ventilation setting was delayed by at least 24 hours compared to the viral shedding of infected recipients from the low ventilation setting. Virus loads in oral swabs followed the same overall trend, although the delay in onset was more subtle compared to nasal washes (Fig. S5). Clinical signs in the high ventilation and low ventilation groups were similar in infected recipients (Fig. S6). Together, our data suggest that in play-based settings with frequent interactions at close-range and multiple transmission modes likely at play, ventilation may not be effective in mitigating influenza virus spread.

**Figure 4.**
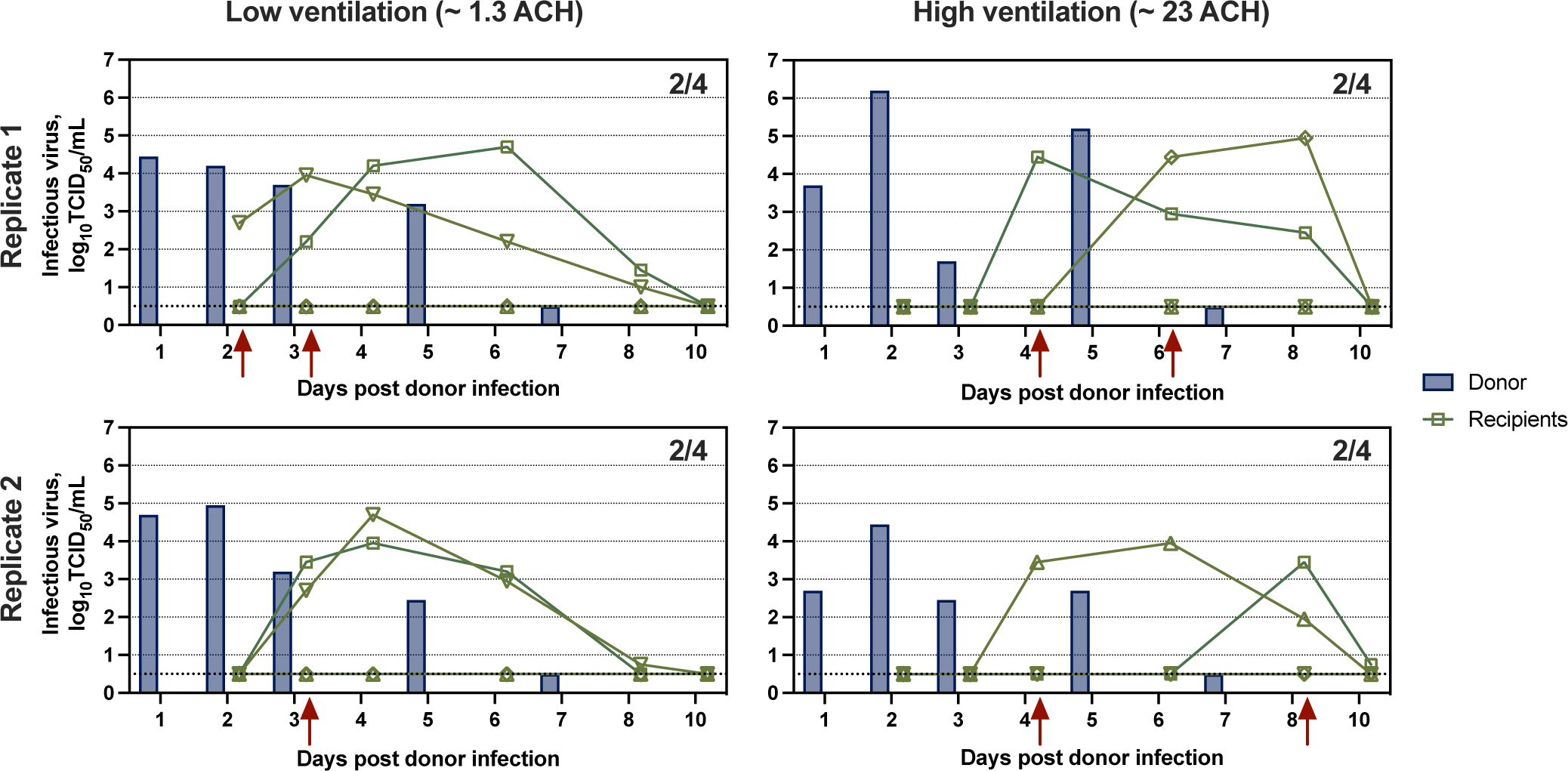
Infectious influenza virus transmission and shedding kinetics in the nasal washes of donor and recipient ferrets under low and high ventilation exposure settings. The transmission exposure occurred the day after inoculation of donors, as indicated in Figure 1A. Dashed lines indicate the limit of detection. Fractions denote the amount of transmission that occurred in each experiment as determined by shedding and seroconversion. Red arrows indicate the days when virus shedding was first detected in nasal washes of recipient ferrets. Donor ferret nasal washes were collected on days 1, 2, 3, 5, and 7 post-infection, and recipient ferret nasal washes were collected on days 2, 3, 4, 6, 8, and 10 post donor-infection. ACH = air changes per hour.

### There is a narrow transmission bottleneck during play-based exposures regardless of ventilation rate

Use of a barcoded virus library enables analysis of not only whether or not transmission occurred, but also how many unique viral genomes established infection in the new host. The barcoded H1N1pdm09 virus has twelve naturally occurring, bi-allelic sites resulting in 4096 unique viral genotypes designed to have equivalent fitness (Fig. S7 and Methods). At 24 hours post-inoculation, nasal washes from donor ferrets exhibited a high amount of viral diversity, evidenced by the detection of > 3,000 barcodes at similar frequencies (Fig. 5). Barcode diversity was maintained at a high level throughout the donor’s infection (Fig. 5). In contrast, the nasal washes of all recipient ferrets that became infected exhibited low barcode diversity, with only one to three prominent barcodes detected in nasal washes (Fig. 5). This narrow bottleneck occurred across both low and high ventilation conditions. Importantly, unique viral barcodes were not repeated across recipients, as expected for fitness neutral genetic variants. These data indicate that ventilation does not appear to impact the viral transmission bottleneck that occurs in the play-based setting.

**Figure 5.**
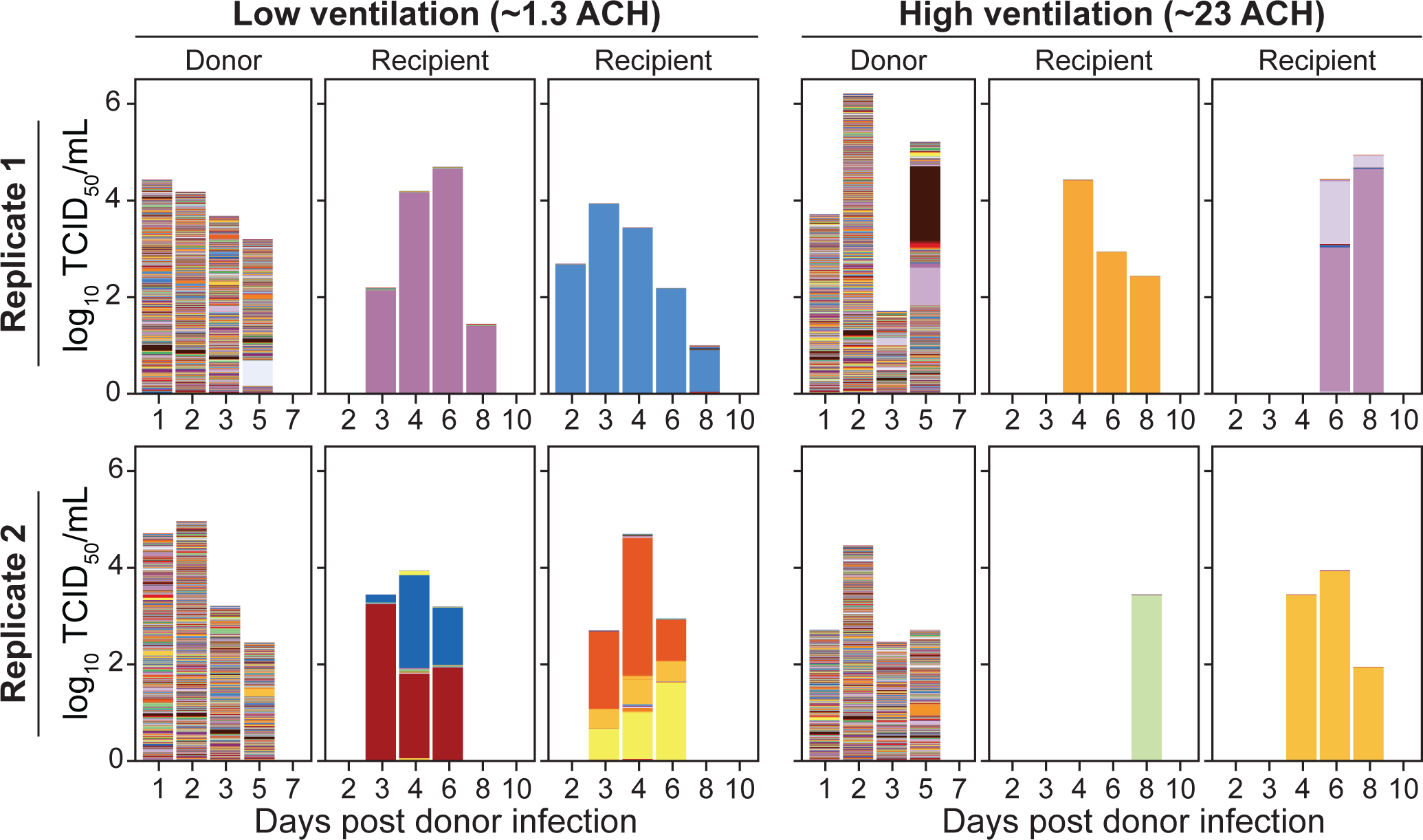
Barcode dynamics in donor and recipient ferrets do not differ according to ventilation Conditions. The height of the bars represents viral titer on a log scale and the colors within each bar show the frequency of individual barcodes within the sample. Samples that had a ≥ 1 log_10_ TCID_50_/mL were analyzed by next-generation sequencing. Each of the 4096 unique barcodes is displayed with a consistent color among all samples. None of the barcodes were shared between recipient ferrets in a singular experiment at frequencies > 1%, emphasizing the neutrality of the barcode system. ACH = air changes per hour.

## Discussion

While the ferret model has been the standard for studying influenza virus transmission dynamics, previous experiments have depended on particular cage configurations to force certain modes of transmission (e.g., airborne using perforated cage divider, all modes by co-housing) and have employed elements (e.g., > 48 hr exposures, directional air flow) (23) that do not reflect the conditions under which humans are typically exposed. Our novel setup aims to simulate more realistic scenarios involving close-range interactions and play, such as those that might occur in a child care setting. This system allows us the opportunity to determine how interventions alter influenza virus spread in a setting with behaviors or interactions that lead to multiple modes of transmission. Here, we focused on ventilation, but future work can build on this system to evaluate whether other engineering controls, such as humidification, directional air flow, filtration, or antiviral coatings on surfaces may more effectively reduce transmission in close-contact scenarios.

Using this modified ferret model, we provide experimental evidence indicating that ventilation in play-based scenarios with frequent interactions at close-range, including with fomites, does not alter transmission efficiency of influenza virus. While many other influenza virus ferret studies use transmission systems with distinct ventilation rates, the direct study of how ventilation influences transmission in the ferret model has not been conducted to date (48). A recent study with guinea pigs found that transmission efficiency was not affected by ventilation airflow speed, and the authors suggested that aerosolized virus resuspended from surfaces played a role in transmission (49). Previous work has drawn correlations between ventilation levels and observed human transmission rates (37, 50), but these studies were not designed to directly assess transmission efficiency derived from ventilation differences.

The effectiveness of an environmental intervention depends not only on the pathogen but also on host factors. The behavior modeled in this experiment is closer to that of children in a child care setting than adults in a workplace. The anatomy and stature of ferrets may contribute to enhanced loss of exhaled respiratory particles by gravitational settling and reduced potential for them to disperse and be removed by ventilation. A 5 μm particle released from a ferret’s nose height, say 5 cm, will settle to the floor in about 1 minute, in contrast to > 20 minutes if released from a young child’s mouth at a height of 100 cm. As acknowledged by the American Society of Heating, Refrigerating and Air-Conditioning Engineers’ (ASHRAE) Standard 241 for control of infectious aerosols, ventilation has limited effectiveness for reducing the risk of transmission in close-range interactions and should be considered as one part of a multi-layered strategy (51). In settings where fewer close-range interactions occur, such as in a school with older children who remain seated at separate desks, office building, or retail store, increasing ventilation could still prove useful in limiting virus transmission.

Our results lend insight to where and how influenza virus transmission may be occurring in settings with close-range interactions. Transmission by direct contact was certainly possible. Environmental sampling provided evidence of influenza virus in aerosols and on surfaces, signifying that both short-range aerosol and indirect contact transmission could be occurring. Surface contamination was similar in both ventilation scenarios, while only slightly less viral RNA was noted in aerosols under increased ventilation. These aerosol viral RNA loads were not as different as would have been expected given the large change in ventilation rate, perhaps due to respiratory particles being removed by gravitational settling, as mentioned above. Most likely, virus concentrations were highest in the immediate vicinity of the ferrets, within a certain radius and low to the ground. Regardless of the underlying reason, NIOSH bioaerosol sampler placement at heights and locations accessible to infected ferrets were meant to represent aerosol exposures likely experienced by recipient ferrets, which ventilation only marginally affected in this study. Additional air samplers may be necessary in future studies to assess how virus-laden aerosols distribute depending on airflow. The modest reduction in viral RNA-containing aerosols in the immediate vicinity of the ferrets was likely insufficient to disrupt transmission by an aerosol route. In addition, transmission by other modes would not be affected by ventilation.

We hypothesized that the low ventilation setting would result in greater viral barcode diversity in recipient nasal washes compared to the high ventilation setting. Contrary to this expectation, the viral populations that established in recipients from both ventilation conditions comprised few barcodes, indicating that ventilation had no major impact on the size of the transmission bottleneck in our system. The observed narrow genetic bottleneck aligns with previous work showing that only a limited number of genotypes establish sustained infection in a recipient (42, 43). These studies showed that, compared to transmission solely through the air, transmission between animals in direct contact yielded a looser genetic bottleneck. While our playpen system is more similar to the direct contact model, the brief exposure times we employed could explain the observed strong constriction in population diversity seen even at low ventilation and with all modes of transmission possible.

Although transmission efficiency was not affected by ventilation rate, shedding kinetics in nasal washes and oral swabs differed. The observed delays in shedding under high ventilation could be due to differences in infectious dose or anatomical site of viral deposition. While preliminary, these findings suggest that different settings or implementation of engineering controls might affect incubation time in situations with close interactions, thus impacting the timing of peak potential for onward transmission. In turn, these effects would carry implications for contact tracing and epidemiological modeling of transmission outbreaks.

There are limitations to the current study. Collection of infectious virus in aerosols from the experimental enclosures during transmission exposures proved difficult despite using three distinct sampling approaches. This outcome aligns with other work, which has also failed to culture influenza virus from aerosols (52, 53), potentially due to dilute aerosol concentrations or harsh sampling techniques. Due to our lack of detection of infectious virus in aerosols, the fate of infectious virus from expelled aerosols under either low or high ventilation remains unclear. Future studies could consider raising sampling flow rates to overcome these technical challenges, although care must be taken to ensure aerosols sampling approaches are not so harsh as to inactivate virus. High flow rates could also increase the effective air exchange rate and affect the spatial distribution of virus in the space. Additionally, while ferrets are a valuable model for studying influenza virus transmission, these animals will never perfectly emulate humans. Ferret behavior may loosely resemble that of small children, but there are critical differences, including the prolonged contact napping of ferrets, as well as extensive biting or chewing, communal eating or drinking, and a lack of vocalization. Detailed analysis of ferret behaviors used during the play-based exposures will shed light on the activities associated with successful influenza virus transmission. Ultimately, human challenge studies or controlled observational epidemiology will be needed to ascertain whether the trends exhibited in this study can be recapitulated among humans. Nonetheless, this work is important to inform real-world settings, and is a critical first step to better characterizing transmission dynamics and evaluating engineering controls in close-contact scenarios where multiple modes of transmission may be operational.

## Materials and Methods

### Influenza virus barcoded plasmid generation

The region of the influenza A/California/07/2009 (H1N1pdm09) genome used for barcode introduction was identified by aligning publicly available sequences of H1N1 viruses circulating in humans. A region with multiple synonymous nucleotide changes was identified from nucleotide position 429 to 489. These naturally occurring mutations were used to build synthetic diversity into the viral stock, thus avoiding the need to introduce foreign sequence into the viral genome (which typically attenuates replication) and limiting the potential for fitness differences among barcoded variants (which would result in their biased replication). Double-stranded Ultramers (IDT) were designed containing degenerate bases with two possible nucleotides at each of the twelve selected barcode sites (Forward: 5’ – GACTCAAGGGGCCTTGCTAAATGACAARCATTCMAATGGRACCATWAARGACAGRAGYCCWTATCGRACYCTAATGAGCTGTCCYATWGGTGAAGTTCCCTCTCCATACAA CTCAAG – 3’; Reverse: 5’ – CTTGAGTTGTATGGAGAGGGAACTTCACCWATRGGACAGCTCATTAGRGTYCGATAWGGRCTYCTGTCYTTWATGGTYCCATTKGAATGYTTGTCATTTAGCAAGGCCCCTTG AGTC – 3’). Carry-over of the NA wild-type sequence was limited by the addition of two stop codons within the barcode region of wild-type plasmid prior to insertion of ultramer DNA. Proper insertion of the ultramers removes the stop codons. Additionally, BarcodeID, our amplicon sequencing analysis software, described below, excludes reads that carry high quality bases that do not match the expected barcode sites. This feature leads to exclusion of any wild-type sequences containing the stop codons from downstream analyses.

Plasmid manipulation for barcode insertion was done as described in Holmes et al. (54), with the exception that the starting material used was the plasmid pHW Cal/07 NA (a kind gift of Dr. Jesse Bloom). Briefly, site-directed mutagenesis was used to introduce an Xho1 site within the wild-type barcode region within the NA segment. Next, the plasmid was linearized with Xho1 digestion, dephosphorylated using rSAP (NEB, Cat No. R0146L), and amplified by PCR using primers that extend outward from the extremes of barcode region (Forward: 5’ – GTGAAGTTCCCTCTCCATACAACTCAAGATTTGAG – 3’; Reverse: 5’ –

CTTGACTCAAGGGGCCTTGCTAAATGAC – 3’). This was followed by PCR purification using the QIAquick PCR Purification Kit (Qiagen, Cat. No. 28106) and elimination of residual wild-type sequences by enzymatic digestion with DPN1 and Xho1. A second PCR purification was performed prior to insertion of the Ultramers into the vector by DNA assembly with the NEBuilder HiFi DNA Assembly Kit (NEB, Cat. No. E2621). Next, the product was transformed into DH5-α cells (NEB, Cat. No. C2987H). Following culture on LB-ampicillin plates (Invitrogen, Cat. No. 45-0034), bacterial colonies were pooled into LB-ampicillin culture media and amplified followed by plasmid purification with the Plasmid Maxi Kit (Qiagen, Cat. No. 12165). Barcode diversity in the plasmid stock was confirmed via next-generation sequencing (Amplicon-EZ – Azenta Life Sciences). The diverse plasmid stock was then used to generate a barcoded H1N1pdm09 virus via reverse genetics by combining it with seven H1N1pdm09 wild-type gene segments encoded in pHW vectors (55).

### Influenza virus and quantification

The barcoded A/California/07/2009 (H1N1; H1N1pdm09) virus was propagated from low multiplicity of infection in MDCK cells (kindly provided by Dr. Daniel Perez, University of Georgia). Infectious virus concentrations were quantified on MDCK cells (kindly provided by Dr. Kanta Subbarao, World Health Organization) using the tissue culture infectious dose 50 (TCID_50_) Spearman Kaber method, as previously described (56).

### Animal work

Male ferrets, aged four to six months, were obtained from Triple F Farms (Sayre, Pennsylvania, USA). Prior to arrival, all ferrets were screened via hemagglutinin inhibition assay for antibodies against circulating influenza A and B viruses, as previously described (57).

All animal work was conducted in a biosafety level 2 at the University of Pittsburgh according to the guidelines of the Institutional Animal Care and Use Committee (approved protocol 22061230). Animals were sedated with isoflurane following approved methods for all nasal washing, oral swabbing, and blood draws.

Donor ferrets were inoculated intranasally with ∼ 10^6^ TCID_50_ of barcoded H1N1pdm09, 250 µL administered into each nostril. Oral swabs and nasal washes were taken for recipient ferrets on days 2, 3, 4, 5, 6, 8, and 10 post donor infection. Donor ferret oral swabs and nasal washes were collected on days 1, 2, 3, 5, and 7 post-infection. Blood draws were collected on days 0, 14, and 21 post donor infection to evaluate seroconversion of donors and recipients, and serological assays were performed as previously described (57). The neutralizing titer was defined as the inverse of the highest serum dilution needed to completely neutralize the infectivity of 10^3.3^ TCID_50_ of virus on MDCK cells.

### Close-range, play-based exposure

The exposure space was comprised of vinyl flooring and a wire cage enclosure intended for use with large animals. The cage enclosure was ∼74 cm tall and had a total footprint of 3.5 m^2^ (Fig. 1). Food, water, and toys were placed in the enclosure. A remote-controlled robot was also deployed, which contained air samplers and sensors as described below under Environmental sample collection. A steel tent frame surrounded the ferret enclosure, and during the low ventilation exposure, a fabric tent cover was attached to the frame and connected to the flooring material with Velcro to minimize air exchange. A large clear plastic viewing window was located on one side of the tent to allow for observation. Three GoPro cameras were placed on the tent frame to record ferret behavior and interactions during the exposure. Aranet4 sensors (SAF, Model TDSPC0H3) were distributed throughout the enclosure and recorded CO_2_, relative humidity, temperature, and partial pressure every minute. One Aranet4 sensor was kept outside of the enclosure as a measure of ambient CO_2_ levels.

Four naïve recipient ferrets and one infected donor ferret (one day post-infection) were placed in the enclosure for four hours. After four hours of continuous exposure, all ferrets were removed and placed in individual housing for the next 10 days.

### Measurement of air exchange rate

To establish the air exchange rate within the tent, the CO_2_ decay method was used (58). Specifically, CO_2_ levels in the tent were increased using dry ice and the atmosphere in the tent was well-mixed using a fan. Once CO_2_ concentrations were sufficiently elevated above the background, the dry ice was removed and the fan was turned off. Aranet4 sensors in the tent recorded CO_2_ every minute. During replicate 1, two sensors were distributed throughout the tent, and during replicate 2, four sensors were distributed throughout the tent. The average CO_2_ concentration measured by all Aranets inside the tent was used for air exchange rate calculations. An Aranet4 sensor was also kept outside the tent as a measure of ambient CO_2_. The air exchange rate was calculated using CO_2_ ratios with the following equation:

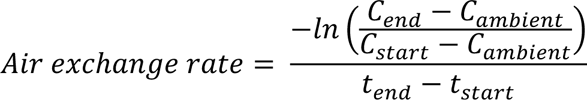

Where C_end_ and C_start_ are the final and initial CO_2_ levels in the tent, respectively, and C_ambient_ is the background level of CO_2_ outside the tent. t_start_ and t_end_ were the times associated with the initial and final CO_2_ concentrations during the decay experiment. Two decay experiments were conducted, prior to each set of replicate exposures, and the average air exchange rate from these replicates was used.

Air exchange rate without the tent was determined using the mechanical ventilation data provided by the building during both experimental replicates. The following equation was used to determine air exchange:

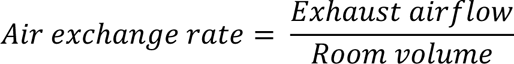

Where exhaust airflow is the rate at which the air is pulled mechanically from the room, in cubic feet per hour, and the room volume is the total volume of the space, in cubic feet. These values were provided by building facilities management. The exhaust airflow was ∼ 1,770 m^3^h^-1^ for the room, which had a volume of 76.5 m^3^, resulting in an air exchange rate of ∼ 23 hr^-1^ for the room.

### Environmental sample collection

Surface samples were collected following each exposure experiment. Sterile foam-tip swabs were used (Puritan, Cat. No. 25-1506 1PF BT) to sample surfaces and subsequently placed in 1 mL collection medium, comprised of 0.5% filter-sterilized bovine serum albumin (Sigma-Aldrich, Cat. No. A3294) in 1X PBS (Sigma-Aldrich, Cat. No. P4417). All swabs were pre-wetted in collection medium. Flat surfaces were swabbed using a 10 cm x 10 cm stencil, and toys, food containers, and water dispensers were swabbed as completely as possible. Three pre-exposure surface samples were taken before each experiment as negative controls. Following collection, surface swab samples were aliquoted for titering and RNA extraction and stored at −80°C until downstream analysis.

NIOSH multi-stage bioaerosol cyclone samplers (Tisch Environmental, Model TE-BC251), a Liquid Spot Sampler aerosol particle collector (Aerosol Devices, Series 110B), and a gelatin cassette were used to collect air samples during each exposure period. NIOSH bioaerosol cyclone samplers were cleaned with isopropanol and milliQ prior to each use. Particles > 4 µm, 1 – 4 µm, and < 1 µm were collected into a sterile 15 mL tube (Falcon, Cat. No. 352096), sterile 1.5 mL tube (Fisherbrand, Cat. No. 02-681-373), and 37 mm polytetrafluoroethylene filter (Millipore, Cat. No. FSLW03700) with backing filter, respectively. Disposable 37 mm sterile plastic cassettes (Pall, Cat. No. 4339-N) were used to collect air on 37 mm sterile gelatin filters (3 µm pore size; Sartorius, Cat. No. 12602-37-ALK) containing a backing filter to ensure the gelatin filter remained intact during sampling. Samplers were connected to GilAir-5 air sampling pumps (Sensidyne, Part 800883-171) with flow rates of 3.5 L/min and 1.5 L/min for the NIOSH bioaerosol cyclone samplers and gelatin cassette, respectively. Two NIOSH bioaerosol cyclone samplers and one gelatin cassette were contained in the robot housing, while the remaining NIOSH bioaerosol cyclone samplers were placed at various locations around the enclosure (Fig. 2). The Spot sampler collected air at ∼ 1.4 L/min with the following settings: 5°C for the conditioner, 40°C for the initiator, 18°C for the moderator, and 27°C for the nozzle. Antistatic silicone tubing (McMaster Carr, Part 1909T39) was used to take in air at ferret height (i.e., 6”) in the close-contact exposure. All samplers ran during the entire exposure period. Baseline air sampling was conducted three to four days prior to each round of experiments. Four of five NIOSH bioaerosol cyclone samplers were processed by adding 500 µL MagMax Lysis/Binding Solution Concentrate (ThermoFisher, Cat. No. AM8500) to each air collection fraction, while the fractions of the fifth NIOSH bioaerosol cyclone sampler were each processed using 1 mL collection medium to assess whether infectious virus could be recovered from NIOSH bioaerosol cyclone samplers. Samples were each vortexed vigorously for one minute, centrifuged briefly, and placed in 1.5 mL microcentrifuge tubes. Gelatin filters were removed from cassettes, placed in 15 mL centrifuge tubes, and dissolved by incubating filters with 1 mL collection medium at 37°C for five minutes, with intermittent vortexing. All samples were stored at −80°C until downstream processing.

### Direct air sampling of donor ferrets

The Spot sampler was used to collect air directly from infected donors on days 1, 2, and 3 post-infection. The collection period was 15 minutes long, resulting in ∼ 21 L of total air sampled. A 7L induction chamber (Vet Equip, Cat. No. 941448) with an inlet and outlet was used to contain the ferret during sampling. Antistatic tubing was connected from the box outlet to the sampler inlet. Makeup air came from the box inlet. Aerosols were collected into 400 µL collection medium, and following sample collection, the total sample volume was increased to 700 µL. To ensure no carry-over of virus, the wick was cleaned between samples by first flushing with 6 mL of isopropanol and subsequently flushing with 18 mL of milliQ water. Clean tubing was used for each sample, and collection vials were cleaned thoroughly with ethanol.

### Infectious virus quantification of environmental samples

Infectious influenza virus concentrations in surface samples and a subset of air samples were determined on Madin-Darby Canine Kidney Cells (MDCK; kindly provided by Dr. Kanta Subbarao) via the Spearman Kaber TCID_50_ assay, as previously described (56).

### Viral RNA quantification of environmental samples

200 µL of each surface sample was extracted using the PureLink Viral RNA/DNA Mini Kit (ThermoFisher, Cat. No. 12280050) according to the standard protocol, with a final elution volume of 50 µL. Air samples were extracted using the QIAamp Viral RNA Mini Kit (QIAGEN, Cat. No. 52904) as specified. 140 µL of Spot samples and gelatin cassette samples were extracted, while 500 µL of NIOSH samples were extracted. All air samples were eluted in 60 µL nuclease-free water. Extracts were stored at −80°C until RNA quantification.

Viral RNA levels in all extracted samples were established via RT-qPCR of the matrix gene, using a previously published set of USDA primers and probe with slight modifications (Forward: 5’-AGATGAGTCTTCTAACCGAGGTCG-3’, Reverse: 5’-GCAAAGACACTTTCCAGTCTCTG-3’, Probe: 5’-FAM-TCAGGCCCCCTCAAAGCCGA-BHQ-3’) (59). All reactions were conducted using the iTaq Universal Probes One-Step RT-qPCR Kit (BioRad, Cat. No. 1725141) on a CFX Connect Real-Time PCR Detection System (BioRad, Cat. No. 1855201) with the following thermocycling conditions: reverse transcription at 50°C for 10 minutes, initial denaturation at 95°C for 2 min, followed by 40 cycles of denaturation and combined annealing/extension at 95°C and 60°C for 10 sec and 20 sec, respectively. In vitro transcribed RNA of the full-length matrix gene was used as the assay standard, and the limit of detection was established as previously described (60). Samples were run in technical duplicates, and any samples that had replicates > 1 Ct apart or had no Ct value for one of two replicates were re-run. If re-runs yielded replicates > 1 Ct apart or had no Ct value for one or both replicates, the viral RNA concentration was defined as below the limit of detection. No template controls were run on each plate.

### Sample processing for next-generation sequencing

Viral RNA extraction from nasal washes of infected ferrets was done using the Quick-RNA Viral 96 Kit (Zymo, Cat. No. R1040) using 160 µL sample volume. This was followed by a one-step RT-PCR reaction using the SuperScript III One-Step RT-PCR System with Platinum Taq High Fidelity DNA Polymerase (ThermoFisher, Cat. No. 12574035) and primers targeting the barcoded region (Forward 5’ – TCGTCGGCAGCGTCAGATGTGTATAAGAGACAGACCTTCTTCTTGACTCAAGGG – 3’; Reverse 5’ – GTCTCGTGGGCTCGGAGATGTGTATAAGAGACAGCAAGCGACTGACTCAAATCTTGA – 3’). The first 33 nucleotides in the primer correspond to adapter sequences for sequencing in the Illumina MiSeq platform. Following amplicon generation, samples were purified using AMPure XP Beads (Beckman Coulter, Cat. No. A63881) with a concentration suitable for the amplicon size (250 bp). Next, DNA quantification was done using the Qubit dsDNA quantification kit (ThermoFisher, Cat. No. Q32851). Samples were submitted to Emory Integrated Genomics Core where indexing (Nextera XT – Illumina), bead cleaning, quality control via bioanalyzer, pooling, and Illumina MiSeq sequencing were completed.

### Amplicon sequencing analysis

Barcode sequences were analyzed using BarcodeID, a software developed in the Lowen laboratory. BarcodeID processes the raw sequences using the BBtools suite (61), before screening reads to ensure that each has the expected base at barcode and non-barcode sites, and that these bases are high quality (Phred values ≥ 30 and ≥ 25, for barcode and non-barcode sites, respectively). Importantly, BarcodeID records the frequency of high-quality bases that don’t match the expected amplicon and barcode sequence, which is used to identify *de novo* mutations that are driving barcode dynamics. No such *de novo* mutations were detected in the present dataset. BarcodeID is available on GitHub at https://github.com/Lowen-Lab/BarcodeID.

## Data, Materials, and Software Availability

All data are available on FigShare upon publication. BarcodeID code is available on Github at https://github.com/Lowen-Lab/BarcodeID.

## Acknowledgments

We thank the MITIGATE FLU team, the Lakdawala Laboratory, and Rachel Duron for critical review and feedback. We thank Milan Shah (Conestoga High School student, Chesterbrook PA) for construction of ERINE. This work was funded by FluLab. Sequencing work was supported by the Emory Integrated Genomics Core Facility (RRID:SCR_023529).

## Notes

### Competing Interest Statement

The authors have declared no competing interest.

