## Supplemental Information for "Ventilation does not affect influenza virus transmission efficiency in a ferret playpen setup"

Current affiliations:
[e]Department of Civil and Environmental Engineering, Duke University, Durham, NC 27708

*Corresponding author
Seema S. Lakdawala


**This PDF file includes:**

Table S1
Figures S1 to S7

**Other supporting materials for this manuscript include:**

Movies S1 and S2

**Table S1.** Seroconversion of donor and recipient ferrets following close-contact, play-based exposure experiments. Titers shown are results of microneutralization assays, details of these are described in Methods.

| Ventilation Condition | Replicate | Ferret | Transmission efficiency | H1N1pdm09 MN titers (Day 0) | H1N1pdm09 MN titers (Day 14) | H1N1pdm09 MN titers (Day 21) |
|---|---|---|---|---|---|---|
| Low | 1 | Donor | | <20 | 5120 | ND |
| | | Recipient | 2/4 | <20, <20, <20, <20 | 7241, <20, <20, 10240 | 6451, <20, <20, 6451 |
| | 2 | Donor | | <20 | 5120 | ND |
| | | Recipient | 2/4 | <20, <20, <20, <20 | 3225, 12902, <20, <20 | 3225, 8127, <20, <20 |
| High | 1 | Donor | | <20 | 6451 | ND |
| | | Recipient | 2/4 | <20, <20, <20, <20 | 5120, <20, 5120, <20 | 3225, <20, 7241, <20 |
| | 2 | Donor | | <20 | 1810 | ND |
| | | Recipient | 2/4 | <20, <20, <20, <20 | 3620, <20, <20, 7241 | 2032, <20, <20, 14482 |

ND = Not determined; MN = microneutralization.

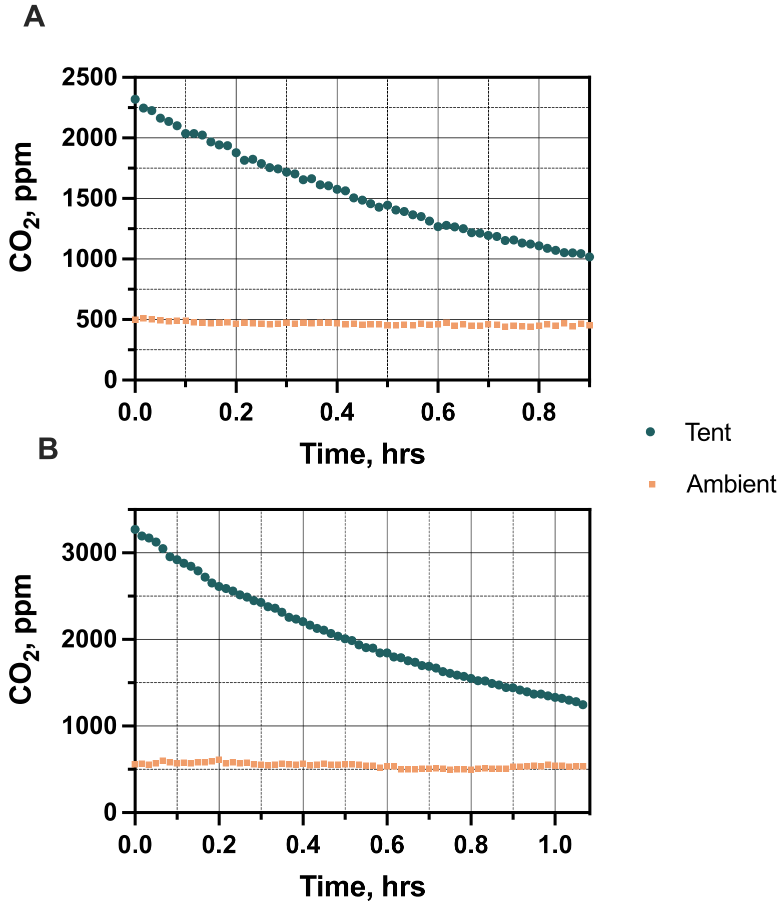


**Fig. S1.** $CO_2$ levels during measurement of air exchange rates in the tent used for low ventilation transmission exposures. (A) Replicate 1 and (B) replicate 2 $CO_2$ decay experiments. Aranet4 sensors were used to measure $CO_2$ measurements of air at multiple locations (n = 2 for replicate 1, n = 4 for replicate 2) within the tent and outside the tent (i.e., ambient). Data was collected every minute. Average $CO_2$ levels within the tent are shown.

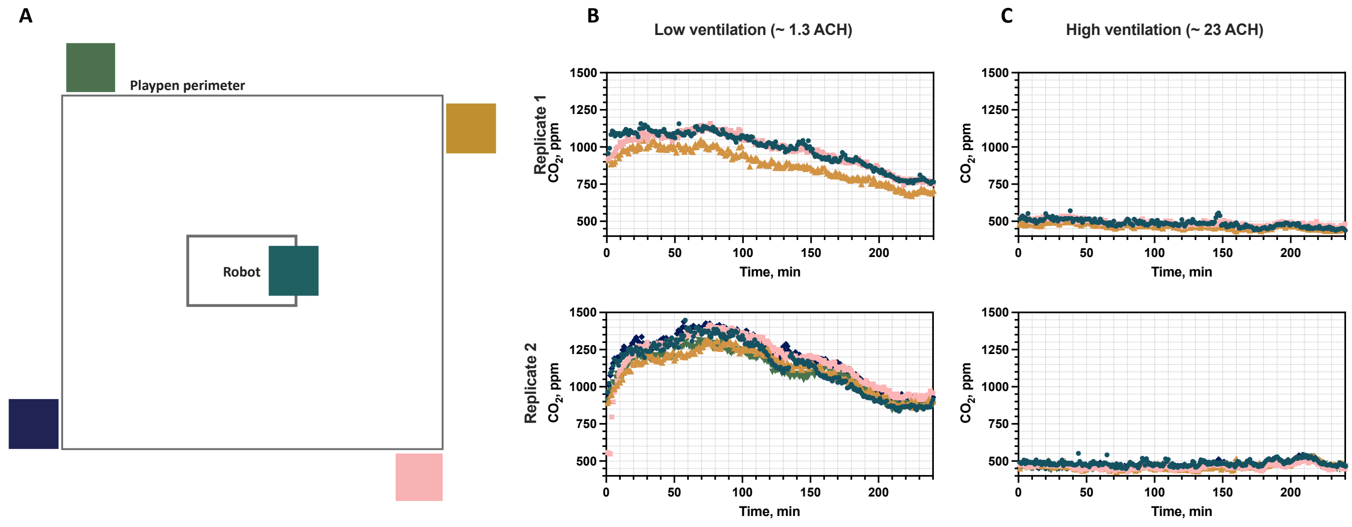


**Fig. S2.** $CO_2$ levels during low and high ventilation transmission exposures. (A) Schematic of $CO_2$ sensor locations and corresponding $CO_2$ levels during the (B) reduced ventilation and (C) increased ventilation exposure settings. The color of symbols in (B) and (C) correspond to the samples of the same color shown in (A). Replicate 1 contains $CO_2$ data from three sensors, while replicate 2 contains $CO_2$ data from five sensors. Sensors collected data every minute.

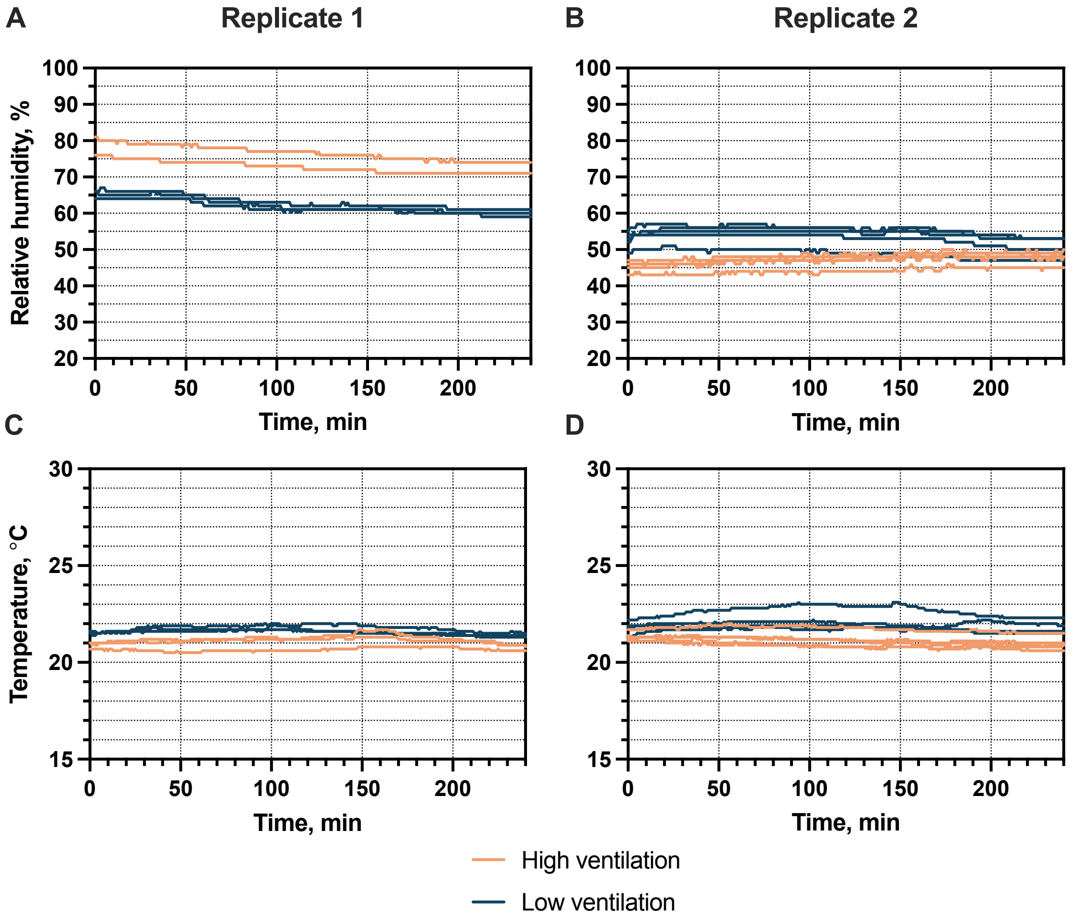


**Fig. S3.** Relative humidity and temperature across experimental replicates of the high and low ventilation transmission exposures. (A) Replicate 1 and (B) replicate 2 relative humidities and corresponding temperatures from (C) replicate 1 and (D) replicate 2.

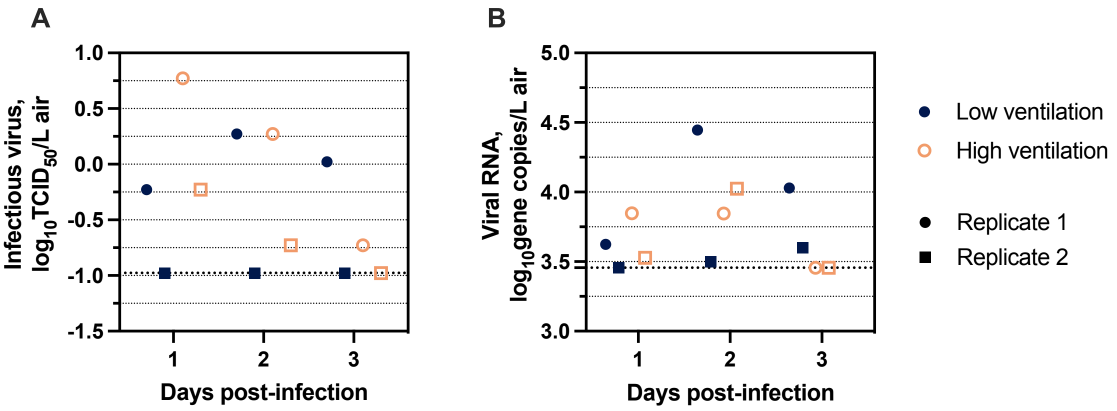


**Fig. S4.** Influenza virus concentrations in the expelled aerosols of infected donor ferrets. (A) Infectious virus and (B) viral RNA concentrations from direct sampling of ferret aerosols using a Spot sampler. Thick dashed lines indicate the limit of detection.

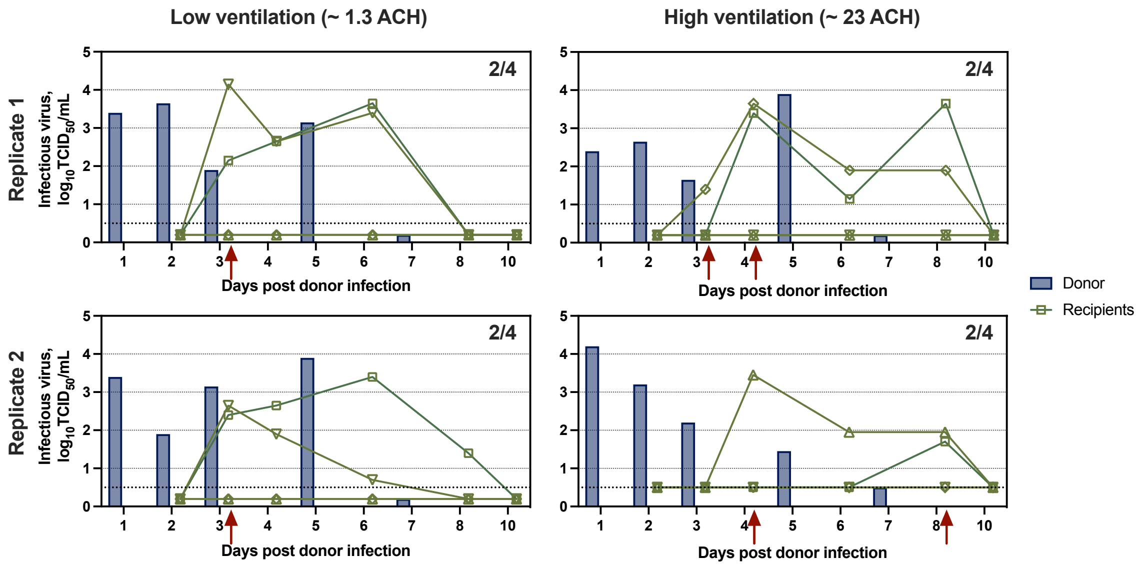


**Fig. S5.** Infectious influenza virus shedding kinetics in the oral swabs of donor and recipient ferrets under low and high ventilation exposure settings. The transmission exposure occurred on day one post donor infection. Dashed lines indicate the limit of detection. Fractions denote the amount of transmission that occurred in each experiment. Red arrows indicate the days when virus shedding was first detected in nasal washes of recipient ferrets. ACH = air changes per hour.

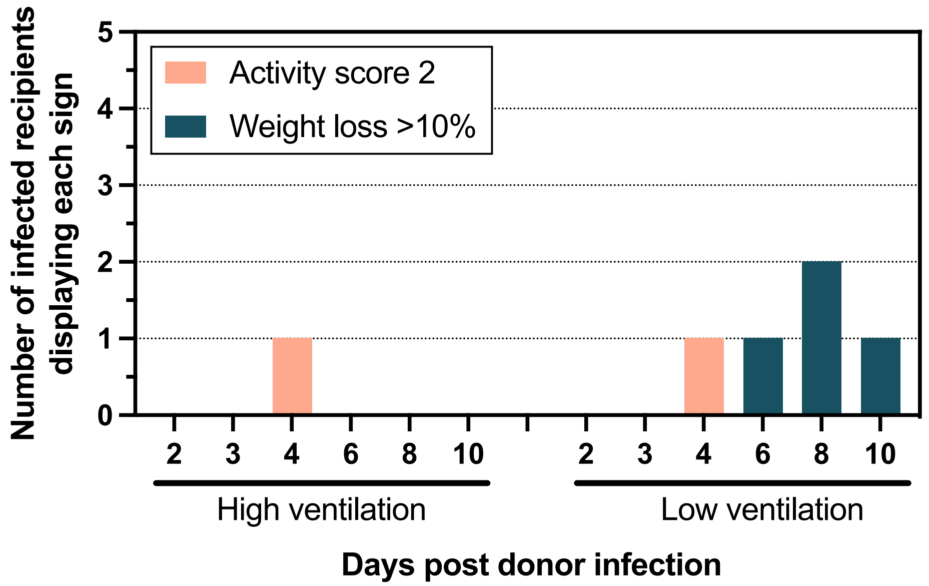


**Figure S6.** Clinical signs of donor and infected recipient ferrets following play-based exposures. Activity scores of 2 indicate the ferret was alert but not at all playful (1).

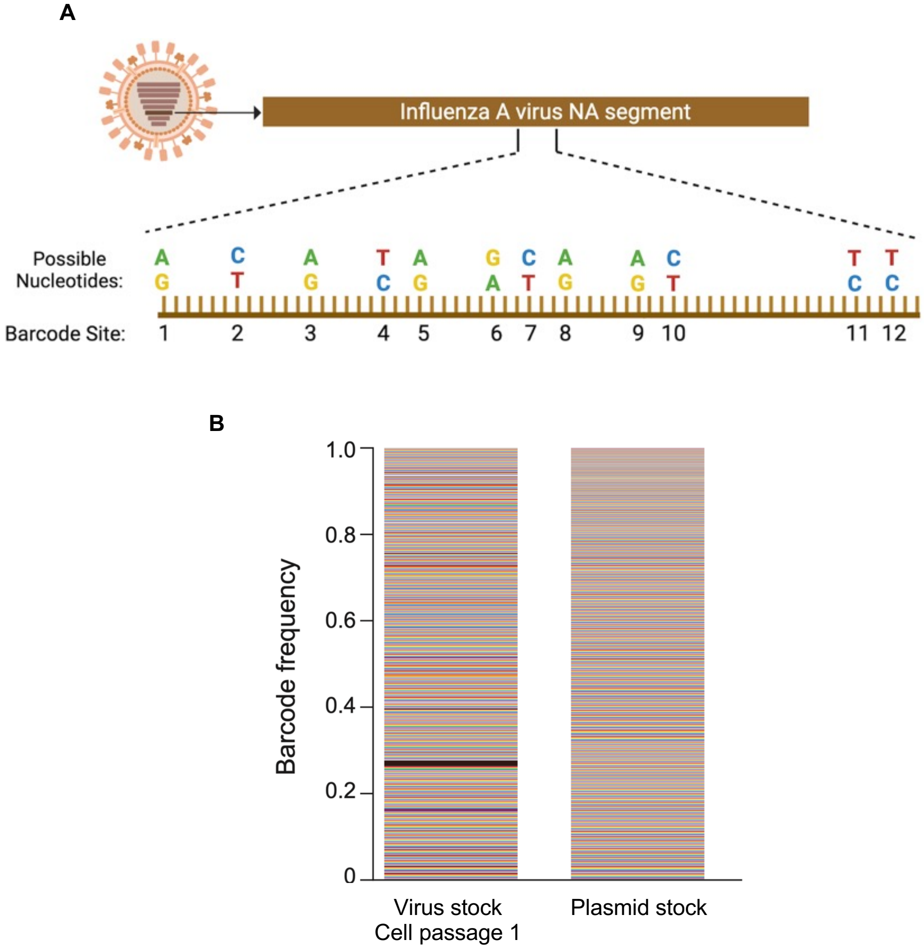


**Figure S7.** Bi-allelic, fitness neutral barcode strategy in the neuraminidase segment produces a highly diverse viral stock. ***(***A) Barcode design for the NA segment of influenza A/California/07/2009 (H1N1pdm09) virus. (B) Barcode diversity in the virus stock (cell passage 1) and the plasmid stock used to generate the virus by reverse genetics. Colors represent unique barcodes, and their frequencies are denoted by the height of the color. Panel A was created in BioRender.com.

**Movie S1 (separate file).** Close-range, play-based exposure under low ventilation conditions.

**Movie S2 (separate file).** Close-range, play-based exposure under high ventilation conditions
